# Synthesis and preclinical evaluation of [^11^C]MTP38 as a novel PET ligand for phosphodiesterase 7 in the brain

**DOI:** 10.1101/2020.10.29.354696

**Authors:** Naoyuki Obokata, Chie Seki, Takeshi Hirata, Jun Maeda, Hideki Ishii, Yuji Nagai, Takehiko Matsumura, Misae Takakuwa, Hajime Fukuda, Takafumi Minamimoto, Kazunori Kawamura, Ming-Rong Zhang, Tatsuo Nakajima, Takeaki Saijo, Makoto Higuchi

## Abstract

**Purpose:** Phosphodiesterase (PDE) 7 is a potential therapeutic target for neurological and inflammatory diseases, although *in-vivo* visualization of PDE7 has not been successful. In this study, we aimed to develop [^11^C]MTP38 as a novel positron emission tomography (PET) ligand for PDE7.

**Methods:** [^11^C]MTP38 was radiosynthesized by ^11^C-cyanation of a bromo precursor with [^11^C]HCN. PET scans of rat and rhesus monkey brains and *in-vitro* autoradiography of brain sections derived from these species were conducted with [^11^C]MTP38. In monkeys, dynamic PET data were analyzed with an arterial input function to calculate the total distribution volume (*V*_T_). The non-displaceable binding potential (*BP*_ND_) in the striatum was also determined by a reference tissue model with cerebellar reference. Finally, striatal occupancy of PDE7 by an inhibitor was calculated in monkeys according to changes in *BP*_ND_.

**Results:** [^11^C]MTP38 was synthesized with radiochemical purity ≥ 99.4% and molar activity of 38.6 ± 12.6 GBq/μmol. Autoradiography revealed high radioactivity in the striatum and its reduction by non-radiolabeled ligands, in contrast with unaltered autoradiographic signals in other regions. *In-vivo* PET after radioligand injection to rats and monkeys demonstrated that radioactivity was rapidly distributed to the brain and intensely accumulated in the striatum relative to the cerebellum. Correspondingly, estimated *V*_T_ values in the monkey striatum and cerebellum were 3.59 and 2.69 mL/cm^3^, respectively. The cerebellar *V*_T_ value was unchanged by pretreatment with unlabeled MTP38. Striatal *BP*_ND_ was reduced in a dose-dependent manner after pretreatment with MTP-X, a PDE7 inhibitor. Relationships between PDE7 occupancy by MTP-X and plasma MTP-X concentration could be described by Hill’s sigmoidal function.

**Conclusion:** We have provided the first successful preclinical demonstration of *in-vivo* PDE7 imaging with a specific PET radioligand. [^11^C]MTP38 is a feasible radioligand for evaluating PDE7 in the brain and is currently being applied to a first-in-human PET study.

## Introduction

Phosphodiesterases (PDEs) are a group of enzymes regulating intracellular levels of cyclic adenosine monophosphate (cAMP) and cyclic guanosine monophosphate (cGMP). These second messengers stimulate intracellular signaling pathways that lead to the activation of protein kinases, resulting in cellular responses in fundamental physiological processes such as neural, cardiovascular, immune, visceromotor, and reproductive functions. PDEs comprise a large superfamily of enzymes that are divided into 11 families [1–3].

PDE7 is composed of PDE7A and PDE7B, both of which specifically hydrolyze cAMP [1–3]. While PDE7A is widely expressed in the brain and peripheral organs [4, 5], PDE7B is abundantly expressed in the brain, particularly in the striatum, nucleus accumbens, olfactory tubercle, thalamus, and hippocampus [5–7]. An increase in intracellular cAMP by PDE7 inhibitors is a potential mechanism of therapeutic approaches to neurological and inflammatory diseases [8]. Indeed, previous studies have demonstrated pharmacologic activities of PDE7-targeting drug candidates in rodent models of Parkinson’s disease [9], Alzheimer’s disease [10], addiction [11], and multiple sclerosis [12].

The roles played by PDE7 in healthy and diseased conditions are likely to be clarified by *in-vivo* visualization of this molecule by positron emission tomography (PET) and specific radioligands. Such technologies also allow measurement of the target occupancy by an exogenous inhibitor for ensuring the target engagement of the agent and for optimizing the therapeutic dosing [13]. Within the PDE families, subtype-selective PET ligands for PDE2A, 4, and 10A [14–16] have been developed. Meanwhile, a Belgian group reported a candidate PET probe for PDE7, [^18^F]MICA-003, with the spiroquinazolinone scaffold as a backbone [17]. However, specific binding of this compound to the targets in living rodent brains could not be proven, since it underwent rapid metabolic conversion to a brain-entering radiomolecule, producing substantive non-specific radiosignals in the brain [17]. Recently, a derivative with the same scaffold was reported to display improved pharmacokinetic properties as examined by a liquid chromatography-tandem mass spectrometry (LC-MS/MS), and autoradiographic labeling of rat brains with a tritiated version of this compound was topologically in agreement with the expression of PDE7B mRNA [18]. <u>As a compound of another group</u>, a series of bicyclic nitrogenated heterocycles was invented as PDE7 ligands by Mitsubishi Tanabe Pharma Corporation and Ube Industries, Ltd. (patent pending: WO 2018/038265, PCT/JP2017/030609). Among these candidates, 8-amino-3-(2S*,5R*-dimethyl-1-piperidyl)-[1,2,4]triazolo[4,3-a]pyrazine-5-carbonitrile (MTP38), was selected with suitability for *in-vivo* imaging of PDE7.

The primary aim of this study was to visualize PDE7 in the brains of rodents and monkeys by PET with [^11^C]MTP38. Furthermore, we determined the occupancy of PDE7 by an inhibitor using [^11^C]MTP38-PET in monkey brains to demonstrate the applicability of this imaging technology to preclinical and clinical evaluations of potential therapeutics acting on PDE7.

## MATERIALS AND METHODS

### Chemistry

MTP38, its radiolabeling precursor 5-bromo-3-(2S*,5R*-dimethyl-1-piperidyl)-8-methoxy-[1,2,4]triazolo[4,3-a]pyrazine (MTP44), MTP-X, and MT-3014 were synthesized at Mitsubishi Tanabe Pharma Corporation.

MTP44 and MTP-X are compounds belonging to the same chemical class as that of MTP38 and are included in the same patent (WO 2018/038265, PCT/JP2017/030609). MTP-X was used as a selective inhibitor of PDE7 in the present work. The Ki value of MTP-X is 10 nM for PDE7 and 1.4 μM or higher for other PDEs. We also employed MT-3014 as a selective inhibitor of PDE10A [19].

### *In-vitro* interactions of MTP38 with PDEs and other bioactive molecules

*In-vitro* inhibitory effects of MTP38 on 12 PDE isozymes (PDE1A, 2A, 3A, 4B1, 5A, 6, 7A, 7B, 8A1, 9A2, 10A2, and 11A4) were determined using a commercial screen panel at Eurofins Panlabs Discovery Services (Taiwan). The enzyme source was bovine retinal rod outer segments for PDE6 or human recombinants for the others. The inhibition percentage at 1 μM or half-maximal inhibitory concentration (IC_50_) value was calculated. All assays were conducted in duplicate. For IC_50_ determination, three independent runs were performed.

The percentage inhibitions of 68 off-target molecules by MTP38 at 1 μM were also assessed using a commercial screen panel at Eurofins Panlabs Discovery Services (Taiwan). This panel included PDE4 but not other PDEs.

### Radiochemistry

A two-step labeling method was employed for the radiosynthesis of [^11^C]MTP38 (Fig. 1). [^11^C]CO_2_ produced by cyclotron (CYPRIS HM-18; Sumitomo Heavy Industries, Tokyo, Japan) was converted to [^11^C]CH_4_, followed by cyanation with ammonium gas over a platinum catalyst to yield [^11^C]HCN [20, 21]. [^11^C]HCN was trapped into a solution of MTP44 (1 mg) and CuI (1 mg) in anhydrous dimethylformamide (0.35 mL), and maintained at 180°C for 5 min to produce intermediate **2**. [^11^C]MTP38 was obtained after nucleophilic substitution of a methoxy group with an amino group using 7N ammonia in methanol (0.3 mL) at 60°C for 1 min. After evaporation with nitrogen gas flow at 100°C for 3 min, 0.5 mL of mobile phase for semi-preparative HPLC (CH_3_CN/0.1 M ammonium acetate, 240:260, vol./vol.) was added, and the mixture was loaded to semi-separative HPLC equipped with a radioactivity detector (column: Inertsil ODS-HL, 10 × 250 mm; column temperature: room temperature; mobile phase: CH_3_CN/0.1 M ammonium acetate, 240:260, vol.vol.; flow rate: 5.0 mL/min; wavelength for ultraviolet detection: 254 nm). The required fraction was collected into a flask to which polysorbate 80 (75 μL) in ethanol (0.3 mL) had been added before radiosynthesis. The solution was then evaporated to dryness, and the residue was dissolved in physiological saline and filtrated through a Millex-GV filter (Merck) to give the final [^11^C]MTP38 formulation for injection.

**Fig. 1.**
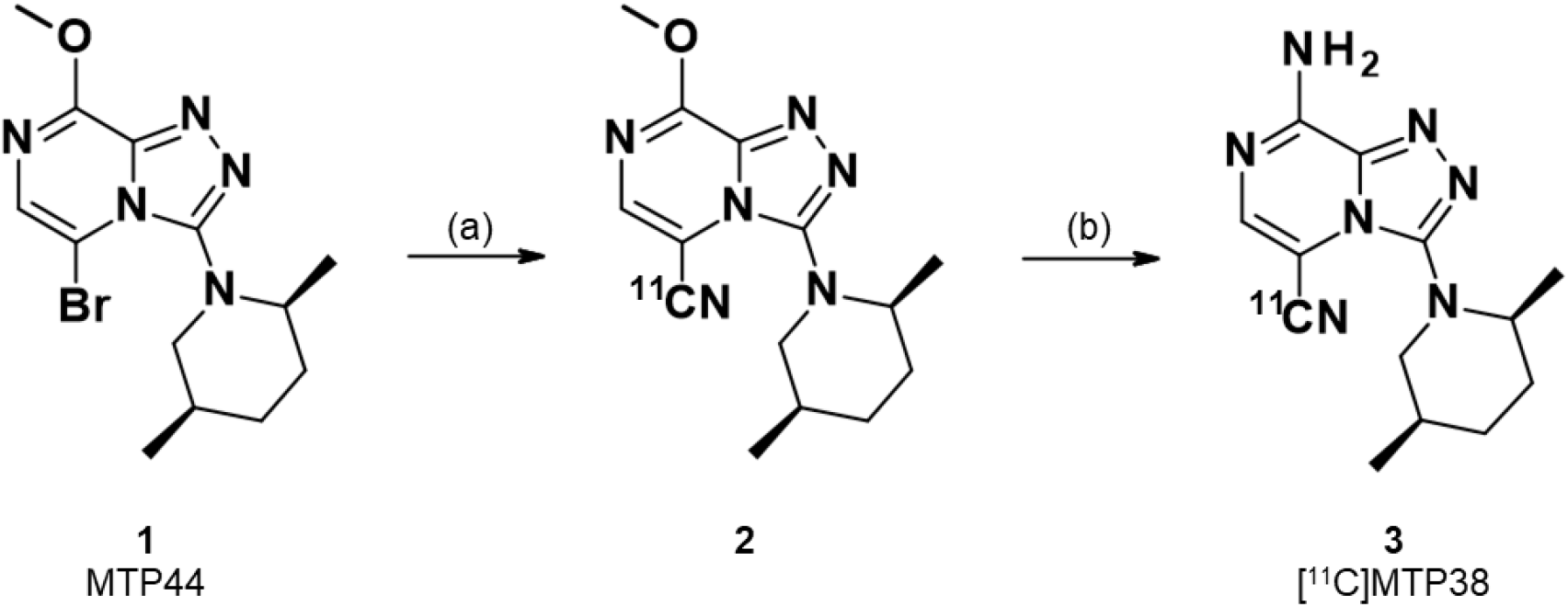
Radiosynthesis of [^11^C]MTP38. (a) [^11^C]HCN, CuI, DMF, 180°C, 5 min; (b) 7N NH_3_ in CH_3_OH, 60°C, 1 min, then 100°C, 3 min, under N_2_ flow.

A portion of [^11^C]MTP38 formulation was also used for analytic HPLC (column: XBridge Shield RP18, 2.5 μm, 3.0×50 mm; mobile phase: 90% CH_3_CN solution/100 mM ammonium phosphate buffer [pH 2.0] and 5 mM 1-octanesulfonic acid sodium salt solution, 40:60, vol.vol.; flow rate: 1.0 mL/min; wavelength for ultraviolet detection: 254 nm) to determine radiochemical purity and molar activity. Radiochemical purity and molar activity of [^11^C]MTP38 were 99.4% or higher and 38.6 ± 12.6 GBq/μmol (mean ± s.d., n=18) at the end of synthesis, respectively.

### *In-vitro* autoradiography of brain slices

*In-vitro* autoradiography was performed using 20-μm-thick fresh-frozen brain sections derived from Sprague-Dawley rats and rhesus monkeys. The sections were preincubated for 10 min in 50 mM Tris-HCl buffer (pH7.4) at room temperature. These samples were then incubated for 60 min at room temperature in fresh buffer containing [^11^C]MTP38 (2 nM) with or without unlabeled MTP38 (10 μM), MTP-X (4 μM), or MT-3014 (40 nM). After the incubation, the brain sections were immersed twice in cold buffer for 3 min, dipped in cold distilled water for 10 sec, and dried with warm airflow. Then the sections were contacted with imaging plates (BAS-MS2025, Fujifilm, Tokyo, Japan) for 60 min. The imaging plates were scanned with an imaging plate reader (BAS5000, Fujifilm, Tokyo, Japan), and the images were analyzed with MultiGuage software (ver. 3.0, Fujifilm, Tokyo, Japan).

### Animals

All animal experiments were carried out in accordance with the Guide for the Care and Use of Laboratory Animals (National Research Council of the US National Academy of Sciences) and were approved by the Committee for the Care and Use of Laboratory Animals of National Institutes for Quantum and Radiological Science and Technology.

Four male Wistar rats (Japan SLC, Inc.) at age 3–5 months were used. These animals were kept in a 12-h dark-light cycle with food and water available *ad libitum*. Two male rhesus monkeys (10 and 12 years, Hamri Co. Ltd, or Japan SLC, Inc.) were also used. The monkeys were kept in individual primate cages in an air-conditioned room. A standard diet, supplementary fruits/vegetables, and a tablet of vitamin C (200 mg) were provided daily.

### Magnetic resonance imaging

Brain structural images were obtained with T2-weighted magnetic resonance imaging using 7.0T Advance system (NIRS/KOBELCO/Bruker) prior to PET experiments. For rats, one of the animals used in the present study was scanned, and the brain image was used as a template. For monkeys, the brains of the two individuals were scanned.

### PET imaging in rats

Each rat was scanned multiple times with a shortest interval of 1 month. Rats were anesthetized with isoflurane (3% for induction and 2% for maintenance) throughout the experiment. The tail vein was cannulated for radioligand injection. Body temperature was maintained at around 37°C using a heating pad. The head of the rat was fixed at the center of the field of view of an animal PET scanner (microPET Focus220, Siemens Medical Solutions, Malvern, USA). A 90-min list-mode data acquisition was started simultaneously with intravenous administration of [^11^C]MTP38. The list-mode data were sorted into 3D sinograms, which were then Fourier-rebinned into 2D sinograms (frames: 10 sec × 6, 30 sec × 8, 1 min × 5, 2 min × 10, and 5 min × 12). The sinograms were reconstructed with a filtered back-projection algorithm using a Hanning filter with a cut-off at the Nyquist frequency (0.5 mm^−1^).

Along with baseline scans, we conducted blockade PET experiments according to the following pretreatment protocols: 1) unlabeled MTP38 (0.5 mg/kg) was administered intravenously at 2 min before radioligand injection, and 2) MTP-X (100 mg/kg) was administered orally at 1 hr before radioligand injection.

### PET data analysis in rats

PET images were co-registered manually onto the magnetic resonance (MR) template image. Details of co-registration were described elsewhere [22]. Regions of interest (ROIs) were placed on the striatum as a target region and on the cerebellum as a reference region according to the expression of PDE7B mRNA in the rat brain [6], and time-activity curves (TACs) were generated for each ROI. Radioactivity in the brain was normalized with injected radioactivity and body weight, and expressed as standardized uptake value (SUV). Then the SUV ratio between the striatum and cerebellum was obtained to estimate the radioligand binding in the target region in baseline and blockade experiments.

### PET imaging in monkeys

Each monkey underwent multiple PET scans with intervals longer than two weeks. The monkeys were given an intramuscular injection of 0.04 mg/kg of atropine sulfate at 1.5 hr before radioligand injection. They were subsequently anesthetized with an intramuscular injection of ketamine (5 mg/kg) and xylazine (0.5 mg/kg), intubated, and kept anesthetized with 1–2% isoflurane. Heart rate, electrocardiogram, body temperature, blood oxygen saturation level, and end-tidal CO_2_ were monitored during the PET scan. The monkeys were placed on a bed in a supine position, and the head was fixed at the center of the field of view of a microPET scanner Focus220. A transmission scan was performed for about 20 min using a ^68^Ge-^68^Ga point source. A 90-min list-mode data acquisition was started simultaneously with intravenous administration of [^11^C]MTP38 via a saphenous vein cannula. All list-mode data were sorted into 3D sinograms, which were then Fourier-rebinned into 2D sinograms (frames: 10 sec × 6, 30 sec × 8, 1 min × 5, 2 min × 10, and 5 min × 12). The images were corrected for attenuation and reconstructed with a filtered back-projection algorithm using a Hanning filter with a cut-off at the Nyquist frequency (0.5 mm^−1^).

Along with baseline scans, we carried out blocking PET experiments according to the following pretreatment protocols: 1) unlabeled MTP38 was administered as an intravenous bolus (0.075 mg/kg) at 1 hr before radioligand injection followed by continuous infusion at a rate of 0.3 mg/kg/h until the end of the PET scan, and 2) MTP-X (0.3–30 mg/kg) was administered orally at 2 or 6 hr before radioligand injection.

### Acquisition of arterial input function

PET scans for monkeys at baseline and following pretreatment with unlabeled MTP38, and 3, 10, 30 mg/kg of MTP-X were performed along with arterial blood sampling to obtain a metabolite-corrected input function. In total, 22 arterial blood samples were collected manually with heparinized syringes via a saphenous artery contralateral to the radioligand injection. Sampling was performed at 10, 20, 30, 40, 50, 60, 75, 90, 105, 120, 150, 180, and 210 sec and at 4, 5, 10, 20, 30, 45, 60, 75, and 90 min after radioligand injection. Blood samples were then placed on ice and divided into two aliquots, one of which was centrifuged with a refrigerated centrifuge (15,000 × *g*, 3 min, 4°C) to obtain plasma.

Radioactivity in blood and plasma was measured with an auto-gamma counter (wizard 2480, PerkinElmer, Waltham, MA, USA). Plasma samples taken at 2, 5, 10, 30, 60, and 90 min were subjected to radiometabolite analysis. An aliquot of plasma was mixed with the same volume of acetonitrile and centrifuged with a refrigerated centrifuge for deproteinization. An aliquot of supernatant (< 1 mL) was injected into an HPLC system (JASCO, Tokyo, Japan) equipped with a radioactivity detector (column: CAPCELL PAK AQ, 5 μm, 10 × 250 mm; column temperature: room temperature; mobile phase: 45% CH_3_CN; flow rate: 6.0 mL/min). The fraction of the parent [^11^C]MTP38 was calculated as a ratio of the peak area of MTP38 to the total area of peaks found on the radiochromatograms. Then an arterial input function was determined by quantifying the metabolite-corrected arterial plasma radioactivity concentration.

### PET data analysis in monkeys

PET images were co-registered onto the individual MR images. ROIs were placed on the prefrontal and medial frontal cortices, hippocampus, amygdala, striatum (caudate + putamen), nucleus accumbens, thalamus, cerebellum, and brainstem. TACs in the baseline and blockade experiments were generated for each ROI.

Regional TACs were examined by several analytical methods to calculate the regional total distribution volume (*V*_T_), which represents the sum of non-specific and specific radioligand binding. Compartment model (1- or 2-tissue) and Logan’s graphical [23] analyses were employed for describing the radioligand kinetics in the brain. The fractional blood volume was fixed at 0.05. For compartment model analyses, an optimal model was chosen on the basis of Akaike’s information criterion (AIC) and model selection criterion (MSC). For graphical analyses, an equilibration time *t*\* was determined so <u>that by the maximum error from the regression within the linear segment was considered to be</u> 10% for each TAC. Time stability of *V*_T_ was investigated by truncating the data duration stepwise from 90 min to 30 min. The *V*_T_ values with or without pretreatment with a blocker were then compared. We also validated the use of the cerebellum as a reference tissue by this comparison, in addition to the autoradiographic findings, in monkey brain slices. The non-displaceable binding potential (*BP*_ND_) was calculated by the original multilinear reference tissue model (MRTM_O_) [24]. *t*\* was determined in the same manner as Logan’s graphical analysis. *BP*_ND_ derived from MRTMo was compared with indirect *BP*_ND_ estimated with Logan’s graphical analysis with an arterial input function as *V*_T,Striatum_/*V*_T,Cerebellum_−1 to justify the employment of MRTMO for the following assays.

All PET image and kinetic analyses described above were performed with PMOD software (ver. 3.711, PMOD Technologies, LLC, Zürich, Switzerland).

### Measurement of PDE7 occupancy by MTP-X in monkey striatum

The occupancy of PDE7 in the monkey striatum by pre-treated MTP-X was calculated as the percentage reduction in *BP*_ND_ from baseline using the following equation:

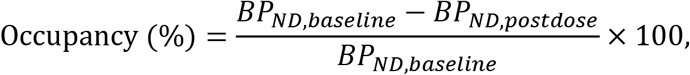

where *BP*_ND,baseline_ and *BP*_ND,postdose_ are the *BP*_ND_ values obtained from PET analyses at baseline condition and after pretreatment with MTP-X, respectively.

The plasma concentration of MTP-X inducing 50% target occupancy (EC_50_) was estimated by the following Hill-Langmuir equation with PRISM software (ver. 7.04, GraphPad Software, San Diego, CA, USA):

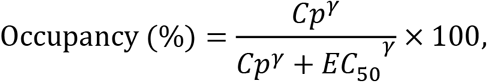

where *C*_p_ is the plasma concentration of MTP-X and γ is the Hill coefficient. The plasma concentration of MTP-X was measured using LC-MS/MS. The method was validated for linearity, precision, accuracy, lower limit of quantification, selectivity, matrix effect, recovery, dilution integrity, carry-over, and stability.

## RESULTS

### *In-vitro* interactions of MTP38 with PDEs and other bioactive molecules

MTP38 potently exerted inhibitory effects on PDE7A and PDE7B with IC_50_ values (nM) of 9.81 (95% confidence interval (CI): 4.45–15.5), 1.21 (95% CI: 0.429–2.05), respectively, while its IC_50_ for PDE2A, 4B1, and 10A2 exceeded 100 nM (Table 1). Moreover, the effects of MTP38 at 1 μM on PDE1A, 3A, 5A, 6, 8A1, 9A2, and 11A4 did not reach 50% inhibition.

**Table 1.** Inhibitory effects of MTP38 on PDEs. Values are geometric mean and 95% confidence interval in three independent experiments. Percent inhibitions of other PDEs (PDE1A, 3A, 5A, 6, 8A1, 9A2, and 11A4) by 1 μM of MTP38 were less than 50%.

| **Isozyme** | **IC<sub>50</sub> (nM)** | **95% CI (nM)** |
| --- | --- | --- |
| **PDE2A** | 114 | 102–127 |
| **PDE4B1** | 269 | 119–429 |
| **PDE7A** | 9.81 | 4.45–15.5 |
| **PDE7B** | 1.21 | 0.429–2.05 |
| **PDE10A2** | 288 | 133–452 |

In a screening panel assay, MTP38 showed no inhibitory effects on the 67 non-PDE molecules tested, with inhibitions at 1 μM being less than 50% (Supplementary Table 1). In line with the above-mentioned PDE assays, MTP38 at 1 μM provoked 74% inhibition of PDE4 (Supplementary Table 1).

### *In-vitro* autoradiography of brain slices

*In-vitro* autoradiograms of rat and monkey brain sections with [^11^C]MTP38 illustrated high radioligand reactivity in the striatum (Fig. 2). While unlabeled MTP38 and MTP-X reduced the striatal radioactivity, MT-3014, a selective PDE10A inhibitor, did not change the radioligand binding. Radioactivity in the cerebellum was relatively low and was not noticeably altered by any of the blockers.

**Fig. 2.**
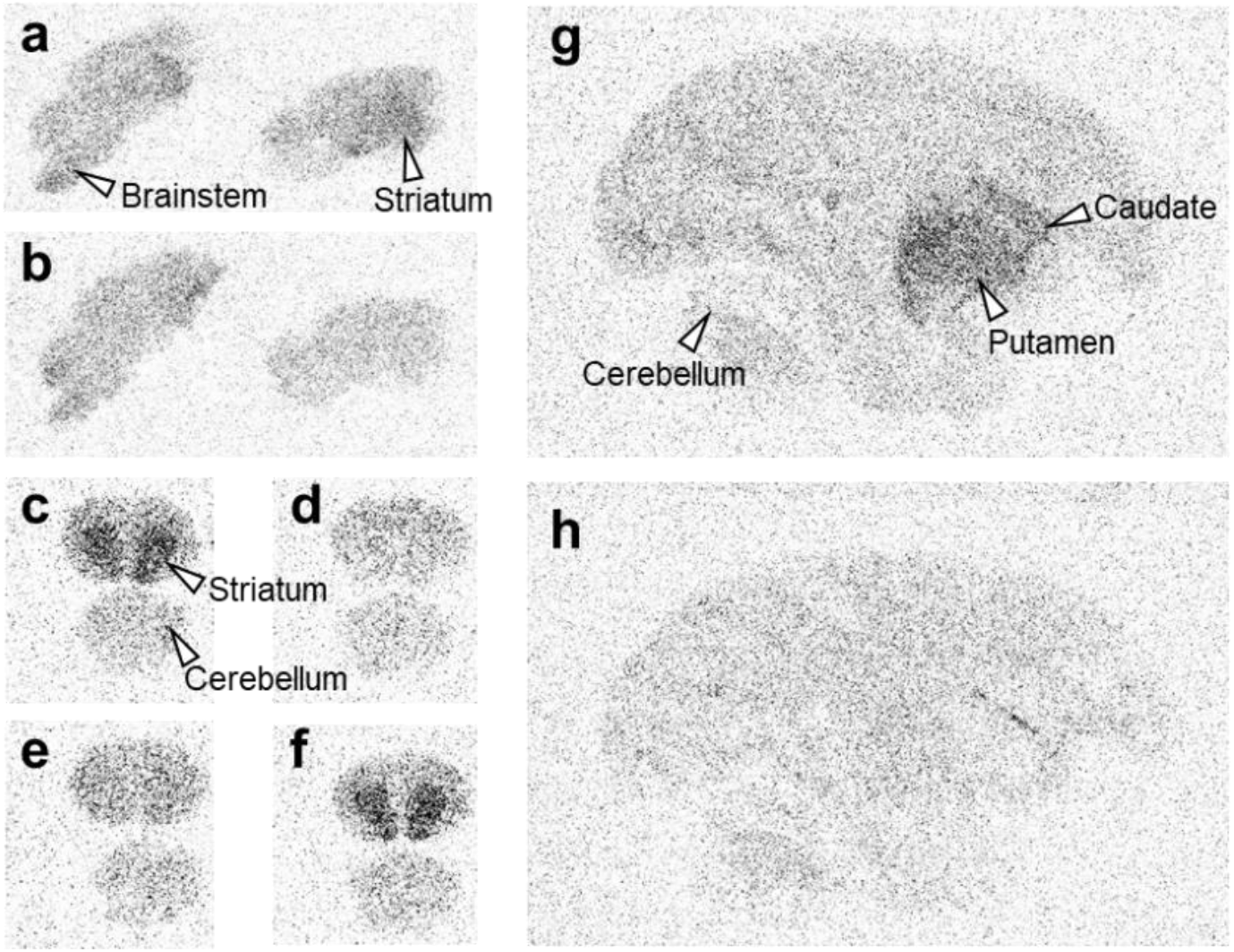
Autoradiographic labeling of rat and monkey brain sections with [^11^C]MTP38. **a, b** Sagittal rat brain sections in the absence (**a**) and presence (**b**) of unlabeled MTP38. **c–f** Coronal rat brain sections in the absence (**c**) and presence of unlabeled MTP38 (**d**), MTP-X (**e**), or MT-3014 (**f**). **g, h** Sagittal rhesus monkey brain sections in the absence (**g**) and presence (**h**) of unlabeled MTP38.

### PET imaging in rats

The radioactivity and molar activity of [^11^C]MTP38 injected into rats were 88–124 MBq and 17.9–37.9 GBq/μmol, respectively. Averaged PET images at 30–60 min after the radioligand injection demonstrated higher radioactivity retention in the striatum than those of other brain regions (Fig. 3b), in accordance with *in-vitro* autoradiographic labeling. In addition to the striatal radiosignals, intensification of radioactivity was observed around the olfactory epithelia (Fig. 3b). Pretreatment of unlabeled MTP38 or MTP-X markedly reduced radioactivity retentions in these areas (Fig. 3c and 3d). TACs indicated that the administered radioligand promptly entered the brain, followed by rapid washout (Fig. 3e and 3f). SUV values in the striatum and cerebellum peaked within 1 min after injection, exceeding 4.0, and declined to a level below 1.0 at 90 min. The striatum-to-cerebellum ratio of SUV plateaued at 5 min and then kept a value approximating 1.5 (Fig. 4). After pretreatment with unlabeled MTP38 or MTP-X, SUV in the striatum was reduced, in contrast to unaltered cerebellar SUV (Fig. 3e and 3f), resulting in a decrease of the SUV ratio to a value around 1.1 (Fig. 4).

**Fig. 3.**
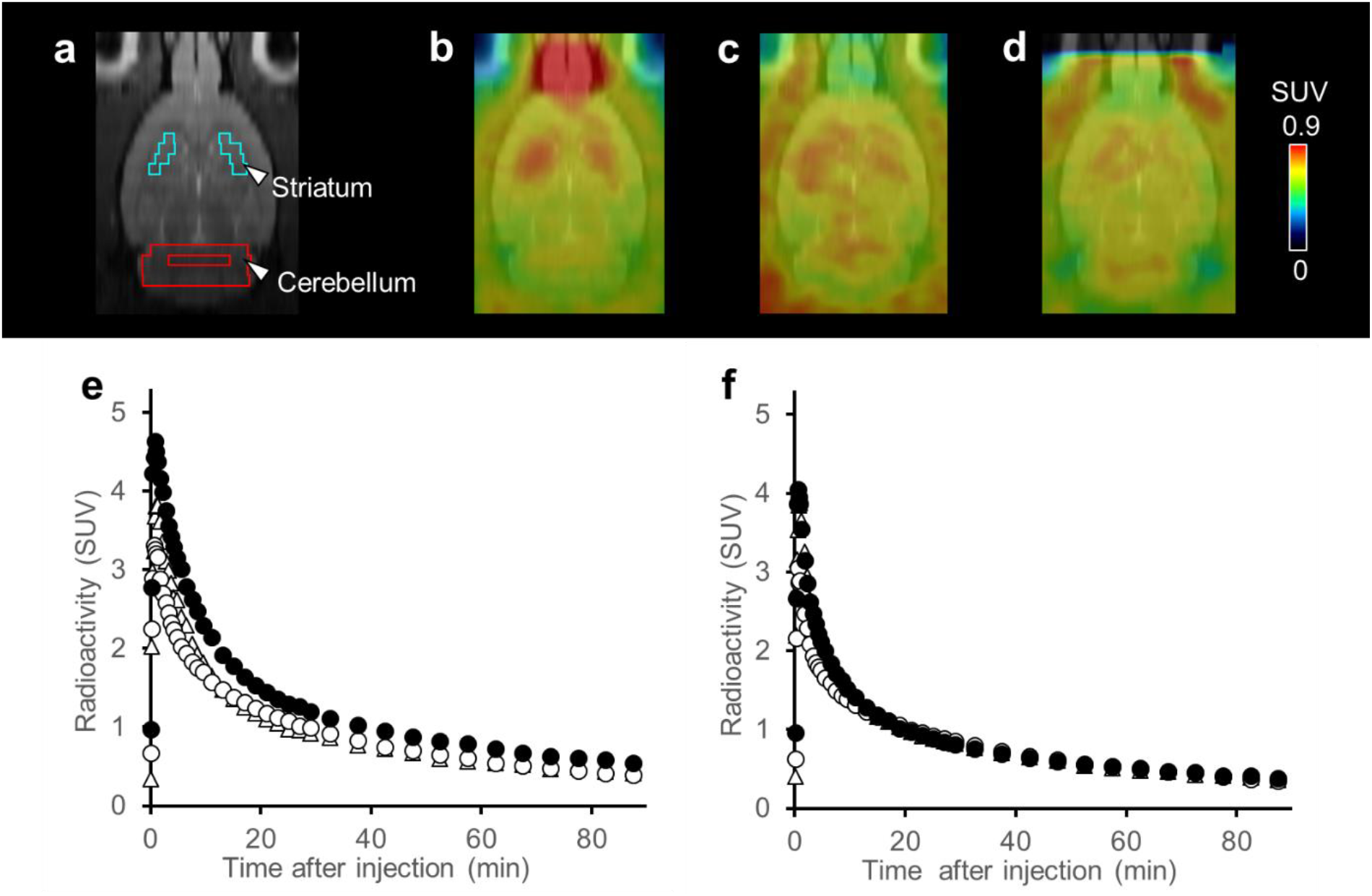
MR, PET images and TACs in the rat brain following injection of [^11^C]MTP38. **a** MR image and ROI settings. **b–d** Representative horizontal PET images averaged at 30–60 min and merged on the corresponding MR template images at baseline (**b**) and following pretreatment with unlabeled MTP38 (**c**) or MTP-X (**d**). **e, f** TACs (means of four animals) in the striatum (**e**) and cerebellum (**f**) at baseline (closed circles), and following pretreatment with unlabeled MTP38 (open circles) and MTP-X (open triangles)

**Fig. 4.**
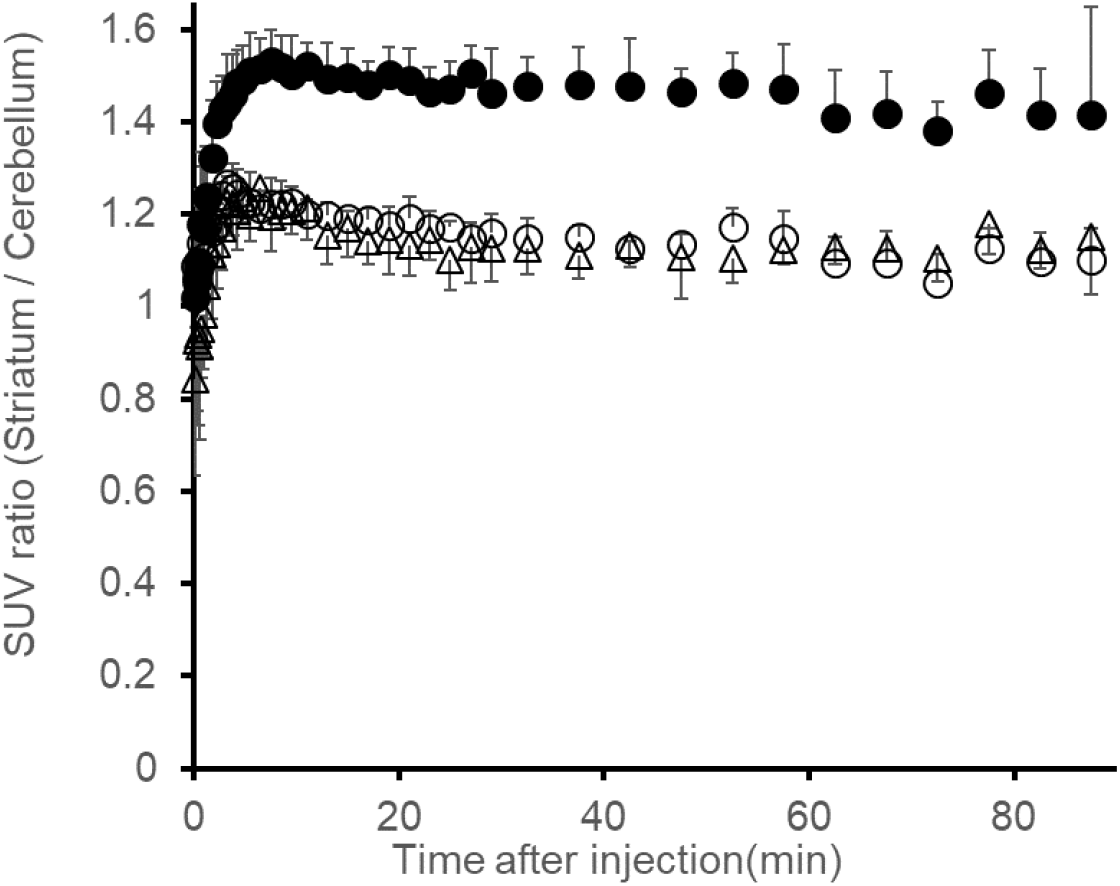
SUV ratio between the striatum and cerebellum in rats. SUV ratio at baseline (closed circles) and following pretreatment with unlabeled MTP38 (open circles) or MTP-X (open triangles). Data are means ± SD of four animals.

### PET imaging in monkeys

The radioactivity and molar activity of [^11^C]MTP38 injected into monkeys were 300–446 MBq and 21.7–29.7 GBq/μmol, respectively. Averaged PET images at 30–60 min showed higher radioactivity retention in the striatum and around white matter than other brain areas under baseline conditions (Fig. 5b). As in rats, a high level of radioactivity was also observed around the olfactory epithelium. The striatal radiosignals were reduced by pretreatment with unlabeled MTP38 (Fig. 5c) or MTP-X (Fig. 5d). TACs indicated that [^11^C]MTP38 rapidly entered the brains and peaked at 3–4 min after injection, with the maximal SUV approximating 5. This was followed by a slightly slower clearance of radioactivity than in rats (Fig. 5d and 5f). As illustrated by PET images, TAC plots demonstrated that the pretreatment with unlabeled MTP38 or MTP-X repressed the striatal radioligand retention (Fig. 5e). By contrast, cerebellar radiosignals were not altered by these pretreatments (Fig. 5f).

**Fig. 5.**
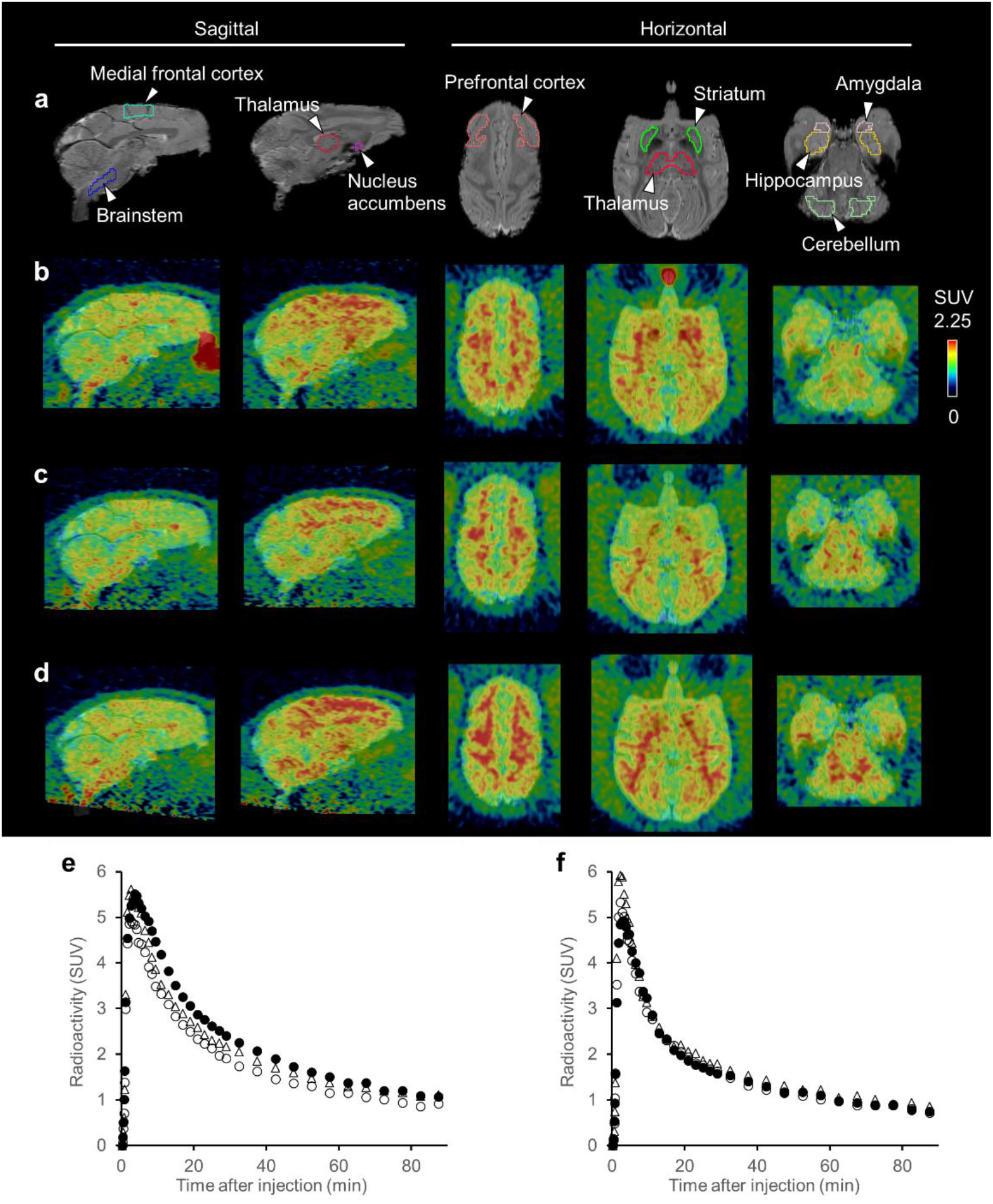
MR, PET images and TACs in the monkey brains following injection of [^11^C]MTP38. **a** MR images and ROI settings. **b–d** Representative (monkey #1) sagittal and horizontal PET images averaged at 30–60 min merged on the individual MR images at baseline (**b**) and following pretreatment with unlabeled MTP38 (**c**) or MTP-X (**d**). **e, f** Representative (monkey #1) TACs in the striatum (**e**) and cerebellum (**f**) at baseline (closed circles) and following pretreatment with unlabeled MTP38 (open circles) and MTP-X (open triangles).

The kinetics of [^11^C]MTP38 in the monkey brain were analyzed with a metabolite-corrected arterial input function (Fig. 6a). The fraction of unmetabolized [^11^C]MTP38 decreased over time, accounting for 27.2% of total radioactivity in plasma at 90 min after injection (Fig. 6b). A major radiometabolite presumably possessing higher polarity than the parent was detected as a peak with a shorter retention time on the radiochromatogram (Fig. 6c).

**Fig. 6.**
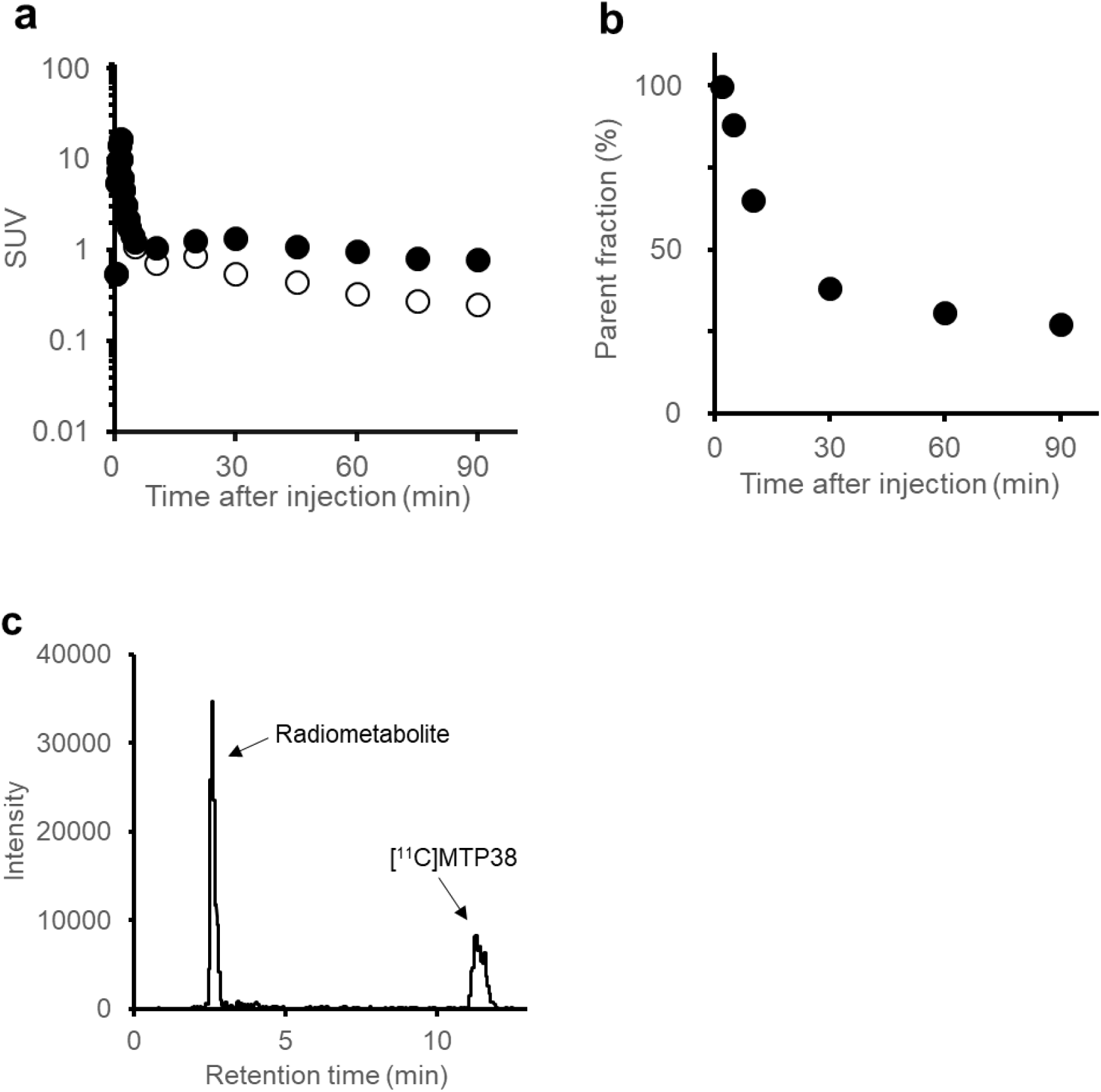
Arterial input function following injection of [^11^C]MTP38 in a monkey. **a** Plasma TAC (closed circles) and radiometabolite-corrected arterial input function (open circles). **b** Time-course changes in the fraction of unmetabolized [^11^C]MTP38 in plasma. **c** Radiochromatogram of plasma at 60 min after [^11^C]MTP38 injection.

*V*_T_ values in defined ROIs were estimated by compartment model analyses and a graphical plot. The 2-tissue compartment model described the radioligand kinetics better than the 1-tissue model based on AIC and MSC (date not shown). However, the 2-tissue compartment model analysis failed to estimate *V*_T_ values in some ROIs including the striatum due to extremely small *k*_3_ values (<0.0001 min^−1^) and consequently unstable *k*_4_ values. On the other hand, plasma-input Logan’s analysis showed linearity of plots after *t*\*=20 min (Supplementary Fig. 1) in all regions. Therefore, *V*_T_ values were determined by plasma-input Logan’s analysis. The highest *V*_T_ value was in the striatum (3.59 mL/cm^3^), followed by the medial frontal cortex (3.27), and the brainstem (3.13) (Table 2). Brain regions displaying low *V*_T_ values included the hippocampus (2.38), amygdala (2.50), and cerebellum (2.69). The *V*_T_ value estimated with 60-min data was 98% of the *V*_T_ value estimated with 90-min data, both in the striatum and cerebellum (Supplementary Fig. 2), justifying the estimation of *V*_T_ values with 90-min dynamic scan data. In addition to the results of *in-vitro* autoradiography, insensitivity of the cerebellar *V*_T_ values to the pretreatments with unlabeled MTP38 and MTP-X supports the view that the cerebellum can be used as reference tissue to quantify the specific binding of [^11^C]MTP38.

**Table 2.** Regional *V*_T_ (mL/cm^3^) of [^11^C]MTP38 in the monkey brain determined by Logan’s graphical analysis

| **Condition<sup>a</sup>** | **STR** | **MFC** | **BS** | **THA** | **NAc** | **PFC** | **CB** | **AMY** | **HIP** |
| --- | --- | --- | --- | --- | --- | --- | --- | --- | --- |
| **Baseline** | 3.59 | 3.27 | 3.13 | 3.03 | 2.97 | 2.93 | 2.69 | 2.50 | 2.38 |
| **Unlabeled MTP38** | 2.70 | 2.98 | 2.95 | 2.79 | 2.39 | 2.56 | 2.51 | 2.16 | 2.05 |
| **MTP-X, 3 mg/kg** | 3.50 | 3.37 | 3.48 | 3.32 | 2.96 | 3.04 | 2.93 | 2.49 | 2.51 |
| **MTP-X, 10 mg/kg** | 3.16 | 3.16 | 3.19 | 2.98 | 2.76 | 2.73 | 2.71 | 2.40 | 2.26 |
| **MTP-X, 30 mg/kg** | 3.08 | 3.11 | 3.21 | 3.05 | 2.70 | 2.78 | 2.78 | 2.40 | 2.26 |

STR, striatum (caudate + putamen); MFC, medial frontal cortex; BS, brainstem; THA, thalamus; NAc, nucleus accumbens; PFC, prefrontal cortex; CB, cerebellum; AMY, amygdala; HIP, hippocampus
Values are means of two animals.
<sup>a</sup>, Baseline or conditions following the administration of indicated blocking agents

*BP*_ND_ values in the striatum were calculated using a graphical analysis based on a reference tissue model, MRTMo (Table 3). *t*\* was 1–3 min. *BP*_ND_ was reduced from 0.409 to 0.157 and from 0.287 to 0.017 in Monkeys #1 and #2, respectively, by the pretreatment with unlabeled MTP38. Pretreatment with MTP-X at multiple doses resulted in a dose-dependent reduction of *BP*_ND_. Finally, *BP*_ND_ estimated by MRTMo was in good agreement with indirect *BP*_ND_, calculated as [striatum-to-cerebellum *V*_T_ ratio–1] (Supplementary Fig. 3). It is considered that *BP*_ND_ calculated with MRTMo can be used to quantify specific radioligand binding.

**Table 3.** Striatal binding potentials of [^11^C]MTP38 in two individual monkeys

| **Animal ID** | **Condition<sup>a</sup>** | **Striatal $BP_{\text{ND}}$** |
| --- | --- | --- |
| **#1** | Baseline | 0.409 |
|  | Unlabeled MTP38 | 0.157 |
|  | MTP-X, 0.3 mg/kg | 0.379 |
|  | MTP-X, 1 mg/kg | 0.369 |
|  | MTP-X, 3 mg/kg | 0.288 |
|  | MTP-X, 10 mg/kg | 0.257 |
|  | MTP-X, 30 mg/kg | 0.191 |
| **#2** | Baseline | 0.287 |
|  | Unlabeled MTP38 | 0.017 |
|  | MTP-X, 0.3 mg/kg | 0.218 |
|  | MTP-X, 1 mg/kg | 0.171 |
|  | MTP-X, 3 mg/kg | 0.115 |
|  | MTP-X, 10 mg/kg | 0.101 |
|  | MTP-X, 30 mg/kg | 0.037 |

<sup>a</sup>, Baseline or condition following the administration of indicated blocking agents

### Measurement of PDE7 occupancy by MTP-X in monkey striatum

Occupancies of PDE7 by MTP-X calculated with *BP*_ND_ values were plotted against the measured plasma concentrations of MTP-X. The plot was then fitted with a direct E_max_ model (Fig. 7). EC_50_ of MTP-X was 688.9 ng/mL, with 95% CI between 390.4 and 1479 ng/mL. The γ and R square values were 0.8323 and 0.7525, respectively.

**Fig. 7.**
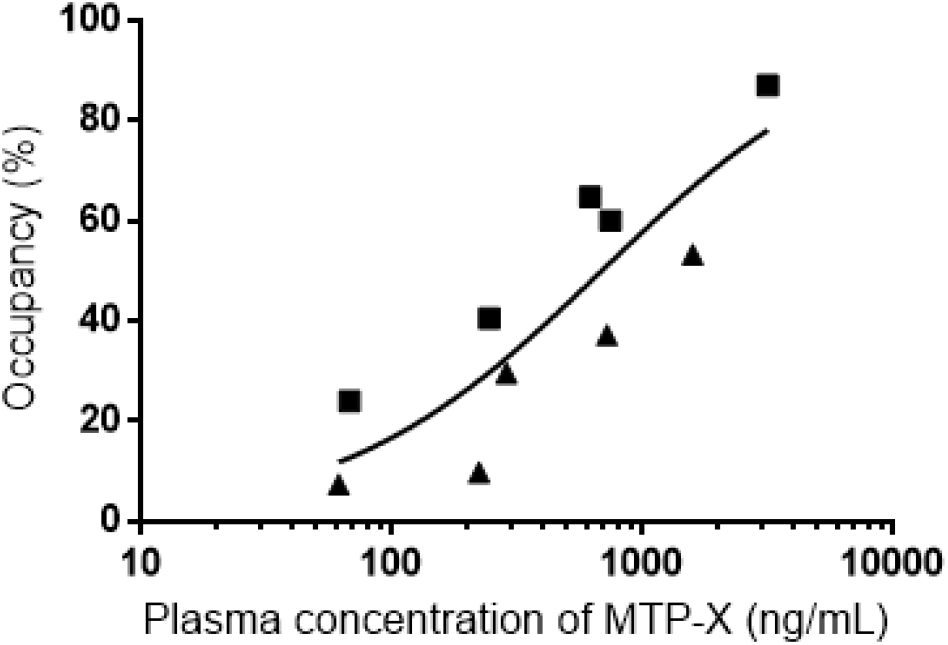
Occupancy of PDE7 by MTP-X. The occupancy of PDE7 by MTP-X in the striatum was plotted against the plasma concentration of MTP-X in two animals (each symbol represents an individual animal). Solid line indicates a sigmoidal curve generated by a fit of the data to a direct E_max_ model.

## DISCUSSION

We developed a novel PDE7-specific PET ligand and applied it to *in-vivo* PET imaging of rat and monkey brains and measurements of PDE7 occupancy by an inhibitor. To our knowledge, this is the first report of successful PET imaging of PDE7 in the brains of living rodents and non-human primates. PET ligands for imaging target molecules with sufficient contrast and accuracy require high affinity and specificity for the binding components. In view of the need for PDE7-acting drugs in the brain without confounding effects on other organs, therapeutic approaches to PDE7B have been of growing interest. PDE7B mRNA expression in the brain is known to be high in the striatum, where other PDEs are also expressed [5, 7]. Therefore, the ligand reactivity with PDE7B and selectivity against other PDEs were considered to be important factors for assessing expressions of PDE7B and evaluating candidate drugs that modify functions of this enzyme. We owned a series of novel compounds with a bicyclic nitrogen heterocycle complex containing [1,2,4]triazolo[4,3-a]piradine as a core structure. Among them, MTP38 was selected as a candidate of a PDE7 PET ligand since it could be labeled with a positron-emitting nuclide, ^11^C, on the cyano moiety and had a high potency for the inhibition of PDE7B (IC_50_. 1.21 nM) and high selectivity for this molecule versus other PDEs. In addition to the ligand affinity for the target molecule (K_d_), the concentration of the binding components could be the major determinant of the target detectability by PET. As a combined measure of the binding affinity and capacity, a B_max_/K_d_ value above 10 in an *in-vitro* binding assay is generally required for the successful application of the potential probe to *in-vivo* PET scans [25]. A recent work reported that the amount of PDE7B protein in the rat brain was 58.0 fmol/mg protein [18]. Provided that the PDE7B protein concentration and IC_50_ of MTP38 are equivalent to B_max_ and K_d_, respectively, the B_max_/K_d_ of this compound is estimated to be 48, which is conceived to be sufficient for the PET detection of PDE7B.

In line with this notion and reported regional levels of PDE7B mRNA expressions [5–7], *in-vitro* autoradiography with [^11^C]MTP38 demonstrated intense radiosignals in the striatum relative to the cerebellum, and this regional difference in autoradiographic labeling was abolished by pretreatment with unlabeled MTP38 and MTP-X, indicating the saturability of [^11^C]MTP38 binding and its specificity for PDE7. In addition, radioligand binding in any of the brain regions was not inhibited by MT-3014, suggesting minimum reactivity of [^11^C]MTP38 with PDE10A.

*In-vivo* PET scans of rats and monkeys revealed favorable kinetics of [^11^C]MTP38 as an imaging agent for PDE7. After intravenous administration, radioactivity was distributed to the brain within a few minutes and then washed out rapidly. In rats, relatively high radioactivity retention was observed in the striatum and around olfactory epithelia. According to the results of *in-vitro* autoradiography, the cerebellum was used as a reference tissue for quantifying specific radioligand binding. The radioactivity concentration ratio of the striatum to the cerebellum at 5–90 min after tracer injection was approximately 1.5 at baseline, and it declined to 1.1 under pretreatment with unlabeled MTP38 or MTP-X. These data indicated the *in-vivo* time-stability and specificity of the striatal [^11^C]MTP38 binding for PDE7 as assayed by the reference tissue model.

In monkeys, the availability of a metabolite-corrected input function acquired by arterial blood samplings enabled detailed kinetic analyses of [^11^C]MTP38 binding in the brains. A single radiometabolite of [^11^C]MTP38 was detected in plasma and was shown to be much more polar than the unmetabolized compound, implying a low probability for this metabolite to penetrate the blood-brain barrier. In fact, *V*_T_ values for [^11^C]MTP38 were well described by analytical models without incorporating the transfer of metabolites from plasma to the brain. This simplicity could provide advantages over a previous PDE7 radioligand, [^18^F]MICA-003 [17], for robust quantitative measurements of target molecules in the brain.

We then compared the accuracy of the methods requiring a plasma input function for estimating parameters of the [^11^C]MTP38 in the brain. The 2-tissue compartment model better described TACs in brain ROIs than the 1-tissue compartment model, but it failed to estimate *V*_T_ values in several regions because it yielded an inadequately small rate constant for the ligand association with targets, *k*_3_. Unlike the compartment model, Logan’s graphical analysis could determine the kinetics in all ROIs examined here (Table 2) and was accordingly chosen as the most reliable method to quantify *V*_T_ values. The linearity of the graphical plot suggested the reversibility of the radioligand binding to PDE7. The time-stability analysis showed that a scan duration of 90 min was sufficient for the quantification of *V*_T_ values, as there was a less than 5% difference between the *V*_T_ values estimated with 60-min and 90-min dynamic data.

We subsequently applied a reference tissue model analysis, MRTMo, for estimation of the radioligand binding without an arterial input function. As in the rat PET assays, the cerebellum was selected as reference tissue since a negligible blocking effect was detected in the autoradiographic experiment. The use of the cerebellum as a reference was also rationalized by the findings that the PDE7 inhibitors did not significantly alter *V*_T_ values in this area (Table 2) and that mRNA expression was noted at the lowest level in the brain [6, 7]. Good agreement was observed between *BP*_ND_ values obtained with MRTMo and indirect *BP*_ND_ values with plasma-input Logan’s analysis (Supplementary Fig. 3; y=1.021x+0.0022, r^2^=0.9987), justifying the application of the reference tissue method for handy measurements of the interaction between [^11^C]MTP38 and PDE7 in the monkey brains.

We accordingly determined the occupancy of striatal PDE7 by the inhibitor, MTP-X, with the use of MRTMo. *BP*_ND_ values in the baseline scans were 0.409 and 0.287 in the two monkeys, and they declined following pretreatment with MTP-X in a manner dependent on the inhibitor dose. The plot of PDE7 occupancy based on the calculation of *BP*_ND_ values versus plasma MTP-X concentration was described by a direct E_max_ model, allowing determination of EC_50_ of MTP-X. A significant portion of the observed data points in the plot rather deviated from the fit curve, presumably due to a low dynamic range of *BP*_ND_ for [^11^C]MTP38. Hence, calculation of the PDE7 occupancy was limited to the striatum most enriched with the target molecules in the brain, and it remains inconclusive whether regional differences in the enzyme occupancy by MTP-X exist.

Despite this shortcoming, our observations support the translational use of [^11^C]MTP38-PET for quantitative assessments of central PDE7 molecules in the human brain under physiological and neuropsychiatric conditions, along with a proof-of-concept clinical study to demonstrate the reactivity of a candidate drug with this enzyme. This neuroimaging platform will also facilitate the development of PDE7-targeting drugs by extrapolating the concentration-occupancy relationship from animals to humans for the optimization of appropriate clinical drug doses.

## CONCLUSION

The current work has provided the first successful *in-vivo* visualization of PDE7 in the brain and has offered an imaging-based methodology for quantifying this molecule and its occupancy by a candidate inhibitor. The radioligand binding to PDE7 is measurable in animal models without the need for arterial blood sampling, and this simplified protocol is potentially implementable in humans. According to the present results and emerging demands on a companion biomarker for clinical trials of potential therapeutics acting on PDE7, clinical studies are ongoing to evaluate the performance of [^11^C]MTP38 in the brains of living individuals.

## ACKNOWLEDGMENTS

We sincerely thank Shoko Uchida, Jun Kamei, Ryuji Yamaguchi, Yuichi Matsuda, and Yoshio Sugii for their assistance in conducting the animal experiments, the staff of the Department of Advanced Nuclear Medicine Sciences, National Institute of Radiological Sciences, for synthesizing the radioligand, and Koji Teshima, Jun Hasegawa, Ryosuke Ide, Rika Nishiyama, and Mitsubishi Tanabe Pharma colleagues for critical discussions of the study plan and results.

## REFERENCES

1. Bender AT, Beavo JA. Cyclic nucleotide phosphodiesterases: molecular regulation to clinical use. Pharmacol Rev. 2006;58:488–520.

2. Keravis T, Lugnier C. Cyclic nucleotide phosphodiesterase (PDE) isozymes as targets of the intracellular signalling network: benefits of PDE inhibitors in various diseases and perspectives for future therapeutic developments. Br J Pharmacol. 2012;165:1288–1305.

3. Azevedo MF, Faucz FR, Bimpaki E, et al. Clinical and molecular genetics of the phosphodiesterases (PDEs). Endocr Rev. 2014;35:195–233.

4. Miro X, Perez-Torres S, Palacios JM, et al. Differential distribution of cAMP-specific phosphodiesterase 7A mRNA in rat brain and peripheral organs. Synapse. 2001;40:201–214.

5. Lakics V, Karran EH, Boess FG. Quantitative comparison of phosphodiesterase mRNA distribution in human brain and peripheral tissues. Neuropharmacology. 2010;59:367–374.

6. Reyes-Irisarri E, Perez-Torres S, Mengod G. Neuronal expression of cAMP-specific phosphodiesterase 7B mRNA in the rat brain. Neuroscience. 2005;132:1173–1185.

7. Kelly MP, Adamowicz W, Bove S, et al. Select 3’,5’-cyclic nucleotide phosphodiesterases exhibit altered expression in the aged rodent brain. Cell Signal. 2014;26:383–397

8. Maliheh S, Maryam B, Mohammad A. New methods for the discovery and synthesis of PDE7 inhibitors as new drugs for neurological and inflammatory disorders. Expert Opin. Drug Discov. 2013;8:733–751.

9. Morales-Garcia JA, Alonso-Gil S, Gil C, et al. Phosphodiesterase 7 inhibition induces dopaminergic neurogenesis in hemiparkinsonian rats. Stem Cells Transl Med. 2015;4:564–575.

10. Bartolome F, de la Cueva M, Pascual C, et al. Amyloid β-induced impairments on mitochondrial dynamics, hippocampal neurogenesis, and memory are restored by phosphodiesterase 7 inhibition. Alzheimers Res Ther. 2018;10:24.

11. Ciccocioppo R, Li H, de Guglielmo G, et al. Phosphodiesterase type 7: a novel target for smoking cessation- preclinical evidence-. Alcohol Alcohol. 2014;49(S1):i34.

12. Martín-Álvarez R, Paúl-Fernández N, Palomo V, et al. A preliminary investigation of phoshodiesterase 7 inhibitor VP3.15 as therapeutic agent for the treatment of experimental autoimmune encephalomyelitis mice. J Chem Neuroanat. 2017;80:27–36.

13. Gunn RN, Rabiner EA. Imaging in central nervous system drug discovery. Semin Nucl Med. 2017;47:89–98.

14. Naganawa M, Waterhouse RN, Nabulsi N, et al. First-in-human assessment of the novel PDE2A PET radiotracer ^18^F-PF-05270430. J Nucl Med. 2016;57:1388–1395.

15. DaSilva JN, Lourenco CM, Meyer JH, et al. Imaging cAMP-specific phosphodiesterase-4 in human brain with *R*-[^11^C]rolipram and positron emission tomography. Eur J Nucl Med Mol Imaging. 2002;29:1680–1683.

16. Boscutti G, Rabiner EA, Plisson C. PET radioligands for imaging of the PDE10A in human: current status. Neurosci Lett. 2019;691:11–17.

17. Thomae D, Servaes S, Vazquez N, et al. Synthesis and preclinical evaluation of an ^18^F labeled PDE7 inhibitor for PET neuroimaging. Nucl Med Biol. 2015;42:975–981.

18. Chen J, Gu G, Chen M, et al. Rapid identification of a novel phosphodiesterase 7B tracer for receptor occupancy studies using LC-MS/MS. Neurochem Int. 2020;137:104735.

19. Koizumi Y, Tanaka Y, Matsumura T, et al. Discovery of a pyrazolo[1,5-a]pyrimidine derivative (MT-3014) as a highly selective PDE10A inhibitor via core structure transformation from the stilbene moiety. Bioorg Med Chem. 2019;27:3440–3450.

20. Oi N, Suzuki M, Terauchi T, et al. Synthesis and evaluation of novel radioligands for positron emission tomography imaging of the orexin-2 receptor. J Med Chem. 2013;56:6371–6385.

21. Iwata R, Ido T, Takahashi T, et al. Optimization of [^11^C]HCN production and no-carrier-added [1-^11^C]amino acid synthesis. Int J Rad Appl Instrum A. 1987;38:97–102.

22. Koga K, Maeda J, Tokunaga M, et al. Development of TASP0410457 (TASP457), a novel dihydroquinolinone derivative as a PET radioligand for central histamine H3 receptors. EJNMMI Res. 2016;6:11.

23. Logan J, Fowler JS, Volkow ND, et al. Graphical analysis of reversible radioligand binding from time-activity measurements applied to [*N*-^11^C-methyl]-(−)-cocaine PET studies in human subjects. J Cereb Blood Flow Metab. 1990;10:740–747.

24. Ichise M, Liow JS, Lu JQ, et al. Linearized reference tissue parametric imaging methods: application to [^11^C]DASB positron emission tomography studies of the serotonin transporter in human brain. J Cereb Blood Flow Metab. 2003;23:1096–1112.

25. Patel S, Gibson R. In vivo site-directed radiotracers: a mini-review. Nucl Med Biol. 2008;35:805–815.

