## Supplementary Table and Figures for "Synthesis and preclinical evaluation of [^11^C]MTP38 as a novel PET ligand for phosphodiesterase 7 in the brain"

**Supplementary Table 1.** Inhibitory effects of MTP38 on various enzymes, ion channels, receptors, and transporters

| Enzymes, ion channels, receptors, and transporters | Sources | Substrates, ligands | Inhibition (%) at 1 μM |
|---|---|---|---|
| Adenosine $A_1$ | Human recombinant | [$^3$H]DPCPX | -12 |
| Adenosine $A_{2A}$ | Human recombinant | [$^3$H]CGS-21680 | 4 |
| Adenosine $A_3$ | Human recombinant | [$^{125}$I]AB-MECA | 10 |
| Adrenergic $\alpha_{1A}$ | Rat submaxillary gland | [$^3$H]Prazosin | 8 |
| Adrenergic $\alpha_{1B}$ | Rat liver | [$^3$H]Prazosin | -3 |
| Adrenergic $\alpha_{1D}$ | Human recombinant | [$^3$H]Prazosin | -3 |
| Adrenergic $\alpha_{2A}$ | Human recombinant | [$^3$H]Rauwolscine | 4 |
| Adrenergic $\beta_1$ | Human recombinant | [$^{125}$I]Cyanopindolol | -2 |
| Adrenergic $\beta_2$ | Human recombinant | [$^3$H]CGP-12177 | -14 |
| Androgen (Testosterone) | Human LNCaP clone | [$^3$H]Methyltrienolone | 1 |
| Bradykinin $B_1$ | Human IMR-90 cells | [$^3$H](Des-Arg$^{10}$)-Kallidin | 9 |
| Bradykinin $B_2$ | Human recombinant | [$^3$H]Bradykinin | 2 |
| Calcium Channel L-Type, Benzothiazepine | Rat brain | [$^3$H]Diltiazem | 16 |
| Calcium Channel L-Type, Dihydropyridine | Rat cerebral cortex | [$^3$H]Nitrendipine | 7 |
| Calcium Channel N-Type | Rat frontal brain | [$^{125}$I]ω-Conotoxin GVIA | 1 |
| Cannabinoid $CB_1$ | Human recombinant | [$^3$H]SR141716A | -6 |
| Dopamine $D_1$ | Human recombinant | [$^3$H]SCH-23390 | -14 |
| Dopamine $D_{2S}$ | Human recombinant | [$^3$H]Spiperone | 4 |
| Dopamine $D_3$ | Human recombinant | [$^3$H]Spiperone | 0 |
| Dopamine $D_{4.2}$ | Human recombinant | [$^3$H]Spiperone | -3 |
| Endothelin $ET_A$ | Human recombinant | [$^{125}$I]Endothelin-1 | -10 |
| Endothelin $ET_B$ | Human recombinant | [$^{125}$I]Endothelin-1 | 3 |

| Enzymes, ion channels, receptors, and transporters | Sources | Substrates, ligands | Inhibition (%) at 1 μM |
|---|---|---|---|
| Epidermal Growth Factor (EGF) | Human A431 cells | [$^{125}$I]EGF | 1 |
| Estrogen ERα | Human recombinant | [$^{3}$H]Estradiol | -1 |
| $GABA_A$, Flunitrazepam, Central | Rat brain (minus cerebellum) | [$^{3}$H]Flunitrazepam | -2 |
| $GABA_A$, Muscimol, Central | Rat brain (minus cerebellum) | [$^{3}$H]Muscimol | -10 |
| $GABA_{B1A}$ | Human recombinant | [$^{3}$H]CGP-54626 | -10 |
| Glucocorticoid | Human recombinant | [$^{3}$H]Dexamethasone | 0 |
| Glutamate, Kainate | Rat brain cortex | [$^{3}$H]Kainic acid | -2 |
| Glutamate, NMDA, Agonism | Rat cerebral cortex | [$^{3}$H]CGP-39653 | -3 |
| Glutamate, NMDA, Glycine | Rat cerebral cortex | [$^{3}$H]MDL 105,519 | -15 |
| Glutamate, NMDA, Phencyclidine | Rat cerebral cortex | [$^{3}$H]TCP | -5 |
| Histamine $H_1$ | Human recombinant | [$^{3}$H]Pyrilamine | -20 |
| Histamine $H_2$ | Human recombinant | [$^{125}$I]Aminopotentidine | -9 |
| Histamine $H_3$ | Human recombinant | [$^{3}$H]N-α-Methylhistamine | 9 |
| Imidazoline $I_2$, Central | Rat cerebral cortex | [$^{3}$H]Idazoxan | -8 |
| Interleukin IL-1 R1 | Human recombinant | [$^{125}$I]Interleukin-1β | -5 |
| Leukotriene, Cysteinyl $CysLT_1$ | Human recombinant | [$^{3}$H]$LTD_4$ | 4 |
| Melatonin $MT_1$ | Human recombinant | [$^{125}$I]2-Iodomelatonin | 7 |
| Muscarinic $M_1$ | Human recombinant | [$^{3}$H]N-Methylscopolamine | -20 |
| Muscarinic $M_2$ | Human recombinant | [$^{3}$H]N-Methylscopolamine | -9 |
| Muscarinic $M_3$ | Human recombinant | [$^{3}$H]N-Methylscopolamine | -21 |
| Neuropeptide Y $Y_1$ | Human SK-N-MC cells | [$^{125}$I]Peptide YY | -13 |
| Neuropeptide Y $Y_2$ | Human KAN-TS cells | [$^{125}$I]Peptide YY | 6 |
| Nicotinic Acetylcholine | Human IMR-32 cells | [$^{125}$I]Epibatidine | -8 |
| Nicotinic Acetylcholine α1, Bungarotoxin | Human RD cells | [$^{125}$I]α-Bungarotoxin | 15 |

| Enzymes, ion channels, receptors, and transporters | Sources | Substrates, ligands | Inhibition (%) at 1 μM |
|---|---|---|---|
| Opiate $\delta_1$ (OP1, DOP) | Human recombinant | $[^3H]$Naltrindole | -4 |
| Opiate κ (OP2, KOP) | Human recombinant | $[^3H]$Diprenorphine | 9 |
| Opiate μ (OP3, MOP) | Human recombinant | $[^3H]$Diprenorphine | 4 |
| Phorbol Ester | Mouse brain | $[^3H]$PDBu | -7 |
| Platelet Activating Factor (PAF) | Human platelets | $[^3H]$PAF | 4 |
| Potassium Channel $[K_{ATP}]$ | Hamster pancreatic HIT-T15 beta cells | $[^3H]$Glyburide | 5 |
| Potassium Channel hERG | Human recombinant | $[^3H]$Astemizole | 17 |
| Prostanoid $EP_4$ | Human recombinant | $[^3H]$Prostaglandin $E_2$ ($PGE_2$) | 8 |
| Purinergic P2X | Rabbit urinary bladder | $[^3H]$α, β-Methylene-ATP | -3 |
| Purinergic P2Y | Rat brain | $[^{35}S]$ATP-αS | -20 |
| PDE4 | Rat brain | $[^3H]$Rolipram | 74 |
| Serotonin (5-Hydroxytryptamine) 5-$HT_{1A}$ | Human recombinant | $[^3H]$8-OH-DPAT | -5 |
| Serotonin (5-Hydroxytryptamine) 5-$HT_{2B}$ | Human recombinant | $[^3H]$Lysergic acid diethylamide (LSD) | -1 |
| Serotonin (5-Hydroxytryptamine) 5-$HT_3$ | Human recombinant | $[^3H]$GR-65630 | -11 |
| Sigma $\sigma_1$ | Human Jurkat cells | $[^3H]$Haloperidol | 6 |
| Sodium Channel, Site 2 | Rat brain | $[^3H]$Batrachotoxinin | 1 |
| Tachykinin $NK_1$ | Human recombinant | $[^3H]$Substance P | 9 |
| Thyroid Hormone | Rat liver | $[^{125}I]$Triiodothyronine | 7 |
| Transporter, Dopamine (DAT) | Human recombinant | $[^{125}I]$RTI-55 | 22 |
| Transporter, GABA | Rat cerebral cortex | $[^3H]$GABA | -2 |
| Transporter, Norepinephrine (NET) | Human recombinant | $[^{125}I]$RTI-55 | 6 |
| Transporter, Serotonin (5-Hydroxytryptamine) (SERT) | Human recombinant | $[^3H]$Paroxetine | -14 |

**Supplementary Fig. 1.** Logan's graphical plot in the monkey brain

Representative (monkey #1) Logan's graphical plots in the striatum (**a**) and cerebellum(**b**).

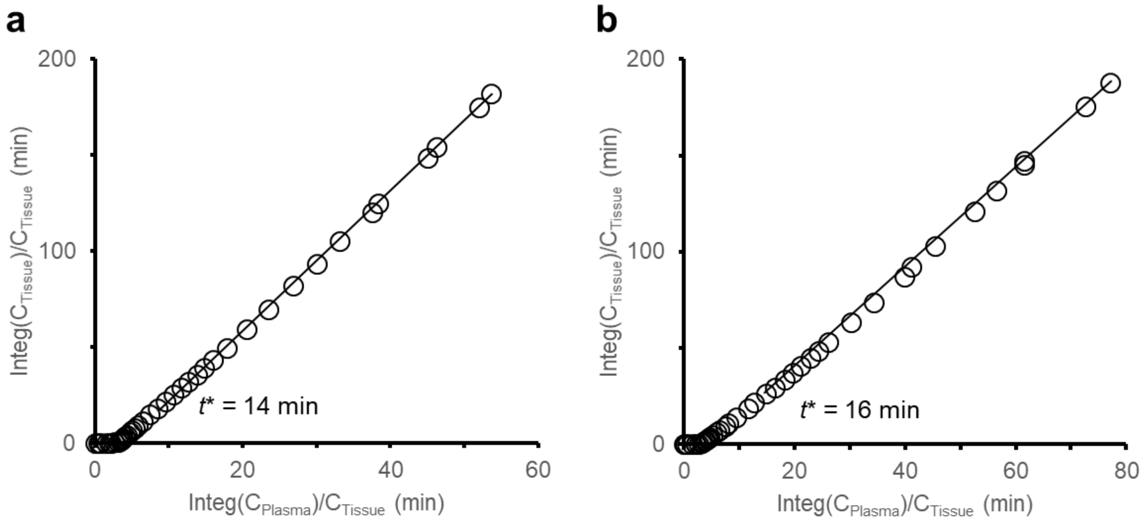

**Supplementary Fig. 2.** Time stability of $V_T$ estimated by Logan's plot in the monkey brain

$V_T$ values in the striatum (circles) and cerebellum (squares) were estimated by Logan's graphical analysis of dynamic PET data and arterial input function truncated from 90 min to 30 min after radioligand injection. Data are means of two animals. Dotted lines indicate 95% and 100% of the $V_T$ value estimated with 90-min data.

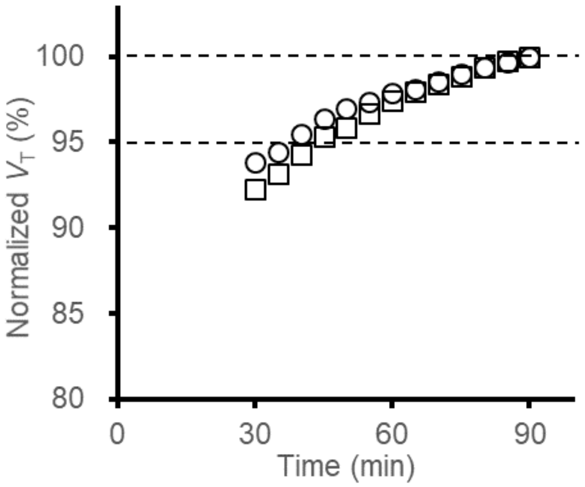

**Supplementary Fig. 3.** Correlations between $BP_{ND}$ values calculated by MRTMo and Logan's plot

Each symbol represents an individual animal. Dotted line indicates the line of identity.

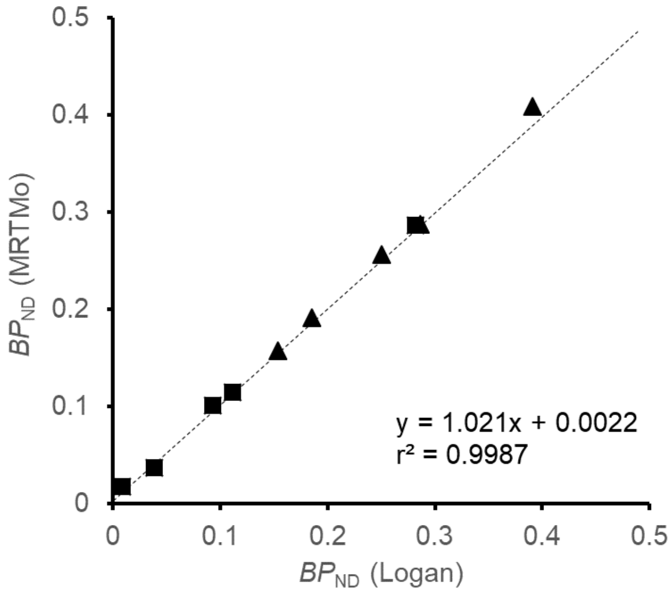
